# Serologic evidence of seasonal influenza A and B viruses in HIV patients on combined antiretroviral therapy in Lagos, Nigeria

**DOI:** 10.1101/553958

**Authors:** AbdulAzeez A. Anjorin, Barakat A. Adepoju

**Author notes:** Corresponding author (AAA).

## Abstract

We investigated serologic evidence of seasonal influenza A and B, and the possibility of their co-infection in HIV patients on combined antiretroviral therapy (cART) in Lagos, Nigeria. A prospective cross-sectional study was designed. A total of 174 HIV positive patients were bled by venipuncture after filling structured questionnaire at the APIN-LUTH clinic, from August to September, 2018. Clear sera were analysed for the detection and quantitative determination of immunoglobulin M specific antibodies to seasonal influenza A subtypes-H1N1 and H3N2, and influenza B by Enzyme Immunoassay (Demeditec, Germany). Results were analysed with Chi-square at 95 % confidence interval. Demographic characteristics showed median age of 44 (mean 45.1, mode 40, range: 18-74) years. Out of the 174 HIV positive patients, 39.7 % (69/174) were seropositive for influenza A and/or B viruses with 58/69 (84.1 %) being positive for influenza A, 11/69 (16 %) for influenza B, and 9/69 (13.4 %) co-infection of influenza A and B. Of the 69 influenza-seropositive patients, age group 41‒ 50 had the highest seroprevalence of 39.1 % (27/69). Females recorded the highest seropositivity of 65.2 % (45/69). Eighty eight (88) % (61/ 69) were on fixed dose cART while 74 % (51/69) were virologically suppressed with HIV RNA < 400 copies/ml. In addition, 2/69 (2.9 %) were positive for HbsAg. Out of the 19/69 (27.5 %) immunocompromised patients (CD4 < 400 cells/mm^3^), 4/19 (21.1 %) were severely immunosuppressed (CD4 < 200 cells/mm^3^). This study revealed serologic evidence of recent circulation of wild influenza A and B viruses in highly suppressed HIV RNA patients on cART. Nonetheless, co-infection with HbsAg and immunocompromised state may further predispose them to serious influenza life threatening complications. Strong advocacy on the need to reduce risk of exposure to influenza, and provision of influenza vaccine in Nigeria are recommended to prevent such complications.

## Introduction

In humans, seasonal influenza virus causes an annual 3-5 million illness and 290,000- 650,000 deaths worldwide (1). Majority of the deaths and severe illness occur in low- and middle-income countries (2) especially Africa with little or no information on epidemiological surveillance. Influenza viral infection can be self-limiting and could have varying severity from being asymptomatic to fatal disease arising from complications. The complications include pneumonia, meningitis and worsening of underlying medical conditions, mostly in immunocompromised individuals (1).

On the other hand, human immunodeficiency virus (HIV) is a retrovirus that spreads through the body rapidly causing damage to the immune system by attacking the CD4 cells with resultant acquired immune deficiency syndrome (AIDS) (3). In 2017, HIV-related causes killed about 1 million people worldwide, having claimed over 35 million lives in the last three decades (4).

According to WHO (2018), newly infected 1.8 million individuals were added to the 36.9 million people living with HIV, with sub-Saharan Africa region having the highest number of 25.7 million infected people (over 66 % of global HIV and more than two thirds of the world's new infections) (4, 5). There are about 197,997,230 people living in Nigeria, equivalent to 2.57 % of the total world population (www.worldometers.info), with up to 4,400,000 people living with HIV (the second-largest HIV epidemic after South Africa and 9 % of total population living with HIV globally), among whom 1,000,000 people are on anti-retroviral therapy (6–8). Lagos State with a total population of about 21 million has HIV prevalence of 2.2 %. Influenza viral infection is a common source of respiratory illness among HIV-infected persons for whom it can be more severe and prolonged (9) while HIV has been described as the most common underlying risk factor in patients having respiratory infection (10). HIV and its comorbidities with influenza virus cause a 4– 8 times higher incidence of hospitalization, and increased death of patients compared to non-HIV infected individuals (11). The reduction of CD4 T-cells due to infection with HIV causes immunodeficiency and thus higher susceptibility to complications of influenza in HIV patients (12). In general, the immunogenicity of HIV infected patients is directly proportional to the CD4 cell count and correlates negatively to the viral load (13). Studies have also shown that better immune health (i.e. higher CD4 cell counts and mostly suppressed viral load of HIV patients on combination antiretroviral therapy) in comparison with immunocompromised patients made them less likely to experience prolonged shedding of influenza virus (9). However, lack of experimental data on seasonal influenza virus in HIV positive individuals is a major problem preventing the development of national policy on preventive strategies in sub-Saharan Africa including Nigeria. There appears to be no study showing co-infection of influenza virus with HIV in Nigeria and that underscores the importance of this study. This study was therefore designed to investigate serologic evidence of seasonal influenza A and B, and the possibility of their co-infection in HIV patients on combined antiretroviral therapy (cART) in Lagos, Nigeria.

## Materials and Methods

### Sample type and study location

Out-patients previously laboratory confirmed to be HIV positive attending AIDS Prevention Initiative in Nigeria (APIN) clinic at the Lagos University Teaching Hospital (LUTH) were recruited for this study from August to September, 2018. APIN-LUTH clinic is a large university-based HIV clinic with about 15,000 patients currently enrolled in Lagos, the largest city in Africa with an estimated population of 21 million. After explaining the concept of the study, a total of 174 unvaccinated HIV patients, age range: 18 to 74 years who gave both oral and written informed consent were bled by venipuncture. Demographic and clinical data were appropriately collected with designed questionnaire. Results of further laboratory analyses including CD4 count, RNA viral load and combined anti-retroviral therapy (ART) regimen were obtained from the patients’ clinical records.

### Ethical Approval and Permission

Permission was sought from the head of APIN clinic where the samples were collected after tendering the ethical approval, reference no: LREC.06/10/1030 obtained from the Health Research and Ethics Committee of the Lagos State University Teaching Hospital (LASUTH). All patients that participated in the study gave both oral and written informed consent.

### Sample collection and Treatment

Approximately 5mL of whole blood samples were collected into sterile plain bottles. They were stored in sample coolers stacked with ice packs before being conveyed to the laboratory. The samples were centrifuged at 3000 rpm for 15 min to obtain clear sera that were aliquoted into labelled sterile plain cryovial tubes. The sera were then stored away at −30°c until ready for serological analysis.

### Laboratory Analysis

Clear sera obtained by centrifugation at 3000 rpm for 15 min were analysed for detection and quantitative determination of IgM antibody specific for Influenza A strains Beijing 265/95(H1N1) and Sydney 5/97(H3N2), and Influenza B strain Hong Kong 5/72 by Enzyme Immunoassay (Demeditec, Germany) in the Department of Microbiology (Virology Research) Laboratory, Lagos State University, Ojo. Following manufacturer’s instructions, sufficient amount of micro titer wells were prepared for the standards, controls and samples as well as for substrate blank. The samples were diluted with ready to use sample diluent provided with the test kit in the ratio 1:101 (2μl serum+200μl sample diluent). 100μl of the ready to use standards, controls and diluted samples were pipetted into the micro titer wells leaving one well empty for the substrate blank. The micro plate was covered with the re-usable plate cover and incubated at room temperature for 60 min. The contents of the wells were emptied and 300μl of diluted washing solution was added. The wells were washed three times while the remaining contents were removed by gently tapping the micro titer plate on a clean tissue cloth. 100 μl of the ready to use enzyme conjugate was added except for the substrate blank. The micro plate was covered before incubation at room temperature for 30 min. The plates were emptied and 300 μl of diluted washing solution was added for washing (x3). Proper draining was ensured. 100 μl of ready to use substrate was added including the substrate blank. The plates were covered and incubated at room temperature in the dark for 20 min. A blue color was observed in the wells before terminating the reaction with 100 μl stop solution, showing a final yellow coloration. Assay absorbance was read at a wavelength of 450nm with micro plate reader (Emax precision, Molecular Devices, California, USA). Sample results were compared with the included standards and controls. They were expressed based on the kit standard curve as recommended by the manufacturer. It should however be noted that positive IgM antibody specific for influenza virus implies recent immunological reaction to circulating live influenza strains as a result of infection since Nigeria does not currently practice influenza vaccination.

### Statistical analysis

Raw data were entered into Microsoft excel, 2013 version. Descriptive statistics was performed while inferential statistics was analysed with Chi-square using GraphPad Prism version 8.0.1 (244), San Diego, USA, putting p-value at < 0.05 statistical significance.

## Results

Demographic characteristics of the patients showed median age of 44, mean 45.1, mode 40, and range: 18-74 years.

Out of the 174 HIV positive patients tested, 69/174 (39.7 %) were seropositive for influenza A and/or B viruses, with 58/69 (84.1 %) positive for influenza A, 11/69 (16 %) for influenza B, and 9/69 (13.4 %) for both influenza A and B (Fig 1).

**Fig 1:**
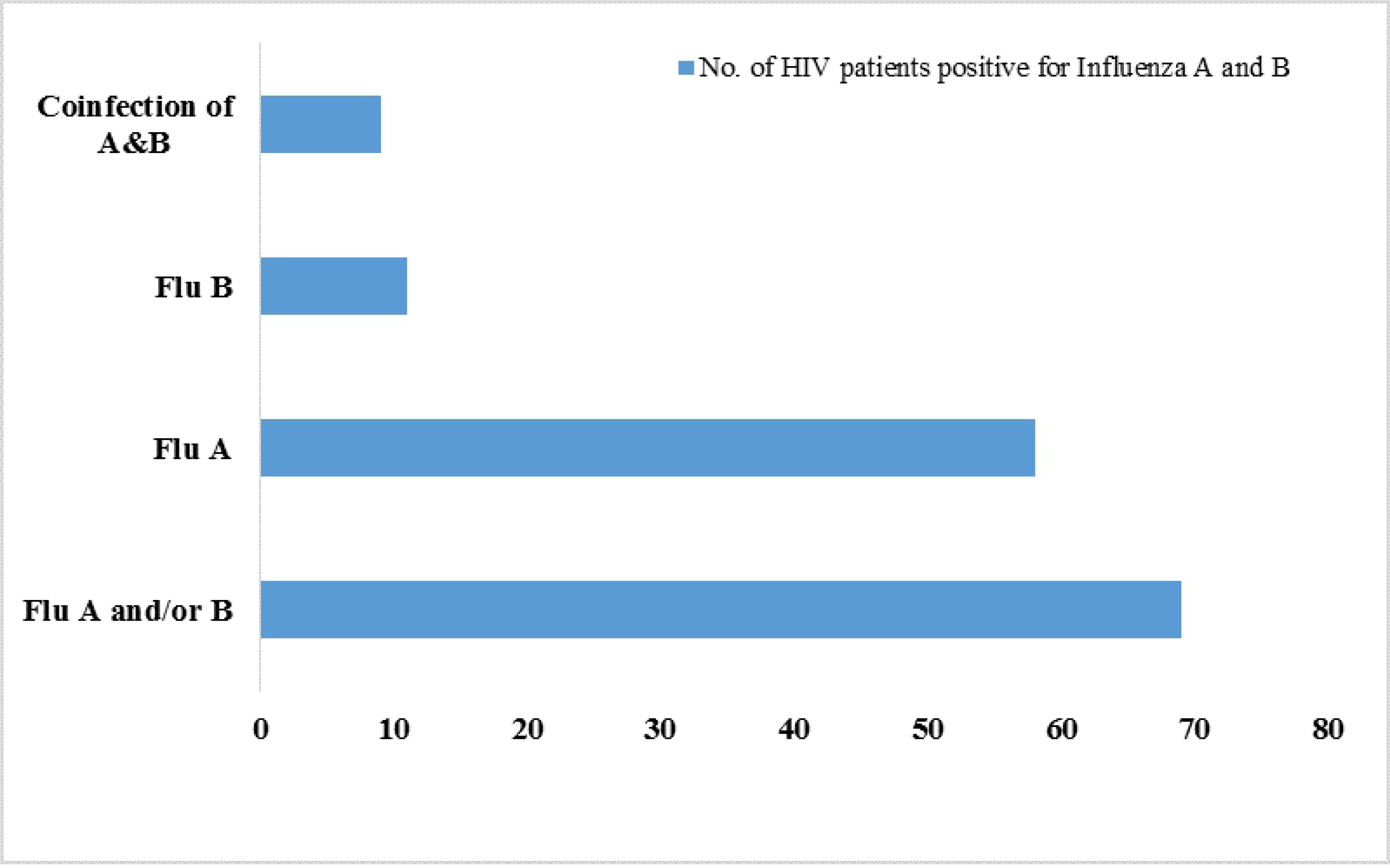
Seroepidemiology of HIV patients positive for Influenza A and B in Lagos, Nigeria.

Age group 41-50 years had the highest seroprevalence of 39.1 % (27/69) with no detection in the age groups 61-70 and 71-80 (Figure 2).

**Figure 2:**
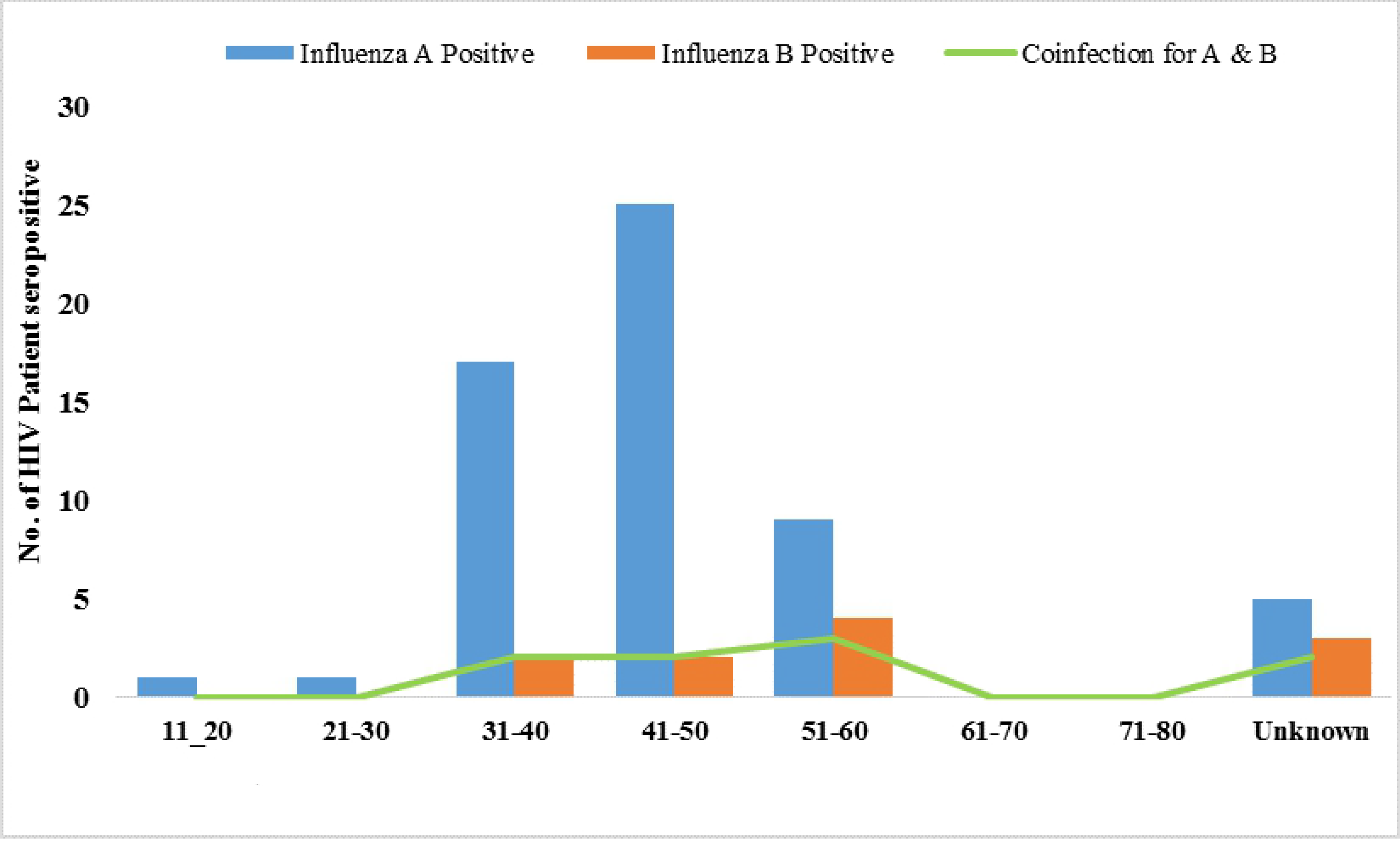
Age distribution of HIV patients seropositive for influenza A and B in Lagos, Nigeria.

Table 1 shows the characteristics of HIV patients tested for influenza A and B viruses. Female patients 45/69 (65.2 %) recorded the highest seropositivity, compare to their male counterpart with 17/69 (24.6 %). Our investigation also revealed that 51/69 (73.9 %) of the patients were virologically suppressed with HIV RNA <400 copies/ mL. Furthermore, a total of 2/69 (2.9 %) patients were positive for HbsAg, whereas 19/69 (27.5 %) were immunocompromised (CD4 <400 cells/mm^3^). Out of the immunocompromised patients, 4/19 (21.1 %) were severely immunosuppressed (CD4 < 200 cells/mm^3^). A total of statistically significant (p=0.0001) 61/ 69 (88 %) HIV patients seropositive to both influenza A and B viruses (compared with seronegative patients) were however found to be on fixed dose combined antiretroviral therapy (cART).

**Table 1:**
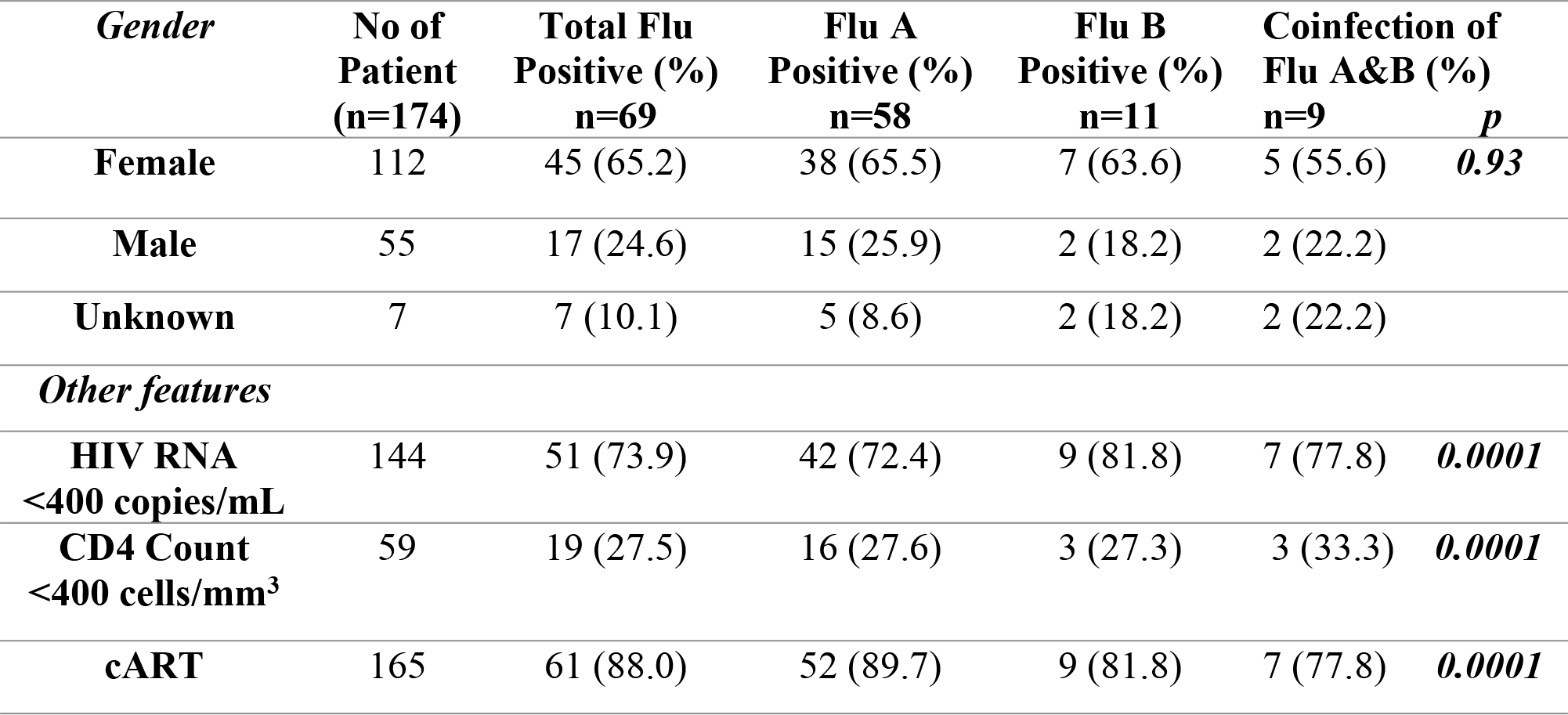
Characteristics of HIV patients positive for Influenza A and B viruses in Lagos, Nigeria.

| **Gender** | **No of Patient (n=174)** | **Total Flu Positive (%) n=69** | **Flu A Positive (%) n=58** | **Flu B Positive (%) n=11** | **Coinfection of Flu A&B (%) n=9** | **p** |
| --- | --- | --- | --- | --- | --- | --- |
| **Female** | 112 | 45 (65.2) | 38 (65.5) | 7 (63.6) | 5 (55.6) | **0.93** |
| **Male** | 55 | 17 (24.6) | 15 (25.9) | 2 (18.2) | 2 (22.2) |  |
| **Unknown** | 7 | 7 (10.1) | 5 (8.6) | 2 (18.2) | 2 (22.2) |  |
| **Other features** |  |  |  |  |  |  |
| **HIV RNA <400 copies/mL** | 144 | 51 (73.9) | 42 (72.4) | 9 (81.8) | 7 (77.8) | **0.0001** |
| **CD4 Count <400 cells/mm<sup>3</sup>** | 59 | 19 (27.5) | 16 (27.6) | 3 (27.3) | 3 (33.3) | **0.0001** |
| **cART** | 165 | 61 (88.0) | 52 (89.7) | 9 (81.8) | 7 (77.8) | **0.0001** |

Table 2 depicted seropositivity to influenza A and B viruses in HIV positive patients on some of the most commonly prescribed combined antiretroviral therapy including: Tenofovir, Emtricitabine, Azidothymidine, Atazanavir, Efavirenz, Nevirapine, Lopinavir/ Ritonavi at the APIN clinic in Lagos.

**Table 2:** Seropositivity of Influenza A and B viruses in HIV patients on most commonly prescribed cART in Lagos, Nigeria.

| **\*ART Regimen** | **No of Patient** | **Total Flu Positive (%)** | **Flu A Positive (%)** | **Flu B Positive (%)** | **Coinfection of Flu A&B (%)** |
| --- | --- | --- | --- | --- | --- |
| FDC(TDF/3TC/EFV) | 74 | 28/74 (37.8) | 23/28 (82) | 5/28 (17.9) | 3/28 (10.7) |
| FDC(AZT/3TC/NVP) | 64 | 20/64 (31.3) | 19/20 (95) | 1/20 (5) | 1/20 (5) |
| FDC(TDF/3TC)-AZT-LPV/r | 4 | 2/4 (50) | 2/2 (100) | 0/2 (0) | 0/2 (0) |
| FDC(TDF/3TC)-LPV/r | 4 | 3/4 (75) | 2/3 (66.7) | 1/3 (33.3) | 1/3 (33.3) |
| FDC(TDF/3TC)-AZT-ATV/r | 3 | 2/3 (66.7) | 1/2 (50) | 1/2 (50) | 1/2 (50) |

\*Shows only the most commonly prescribed cART for HIV patients in Lagos.

## Discussion

Serological assay usually complement epidemiological and clinical investigations in the detection and identification of influenza viruses. They measure antibodies developed in response to infection with antigenically novel and seasonal influenza viruses (14). ELISA method has been used in different studies for detecting serological evidence of influenza virus in humans (15–17) because it is apt, easy, fast, can be automated and commercially available for large-scale serosurvey, targeting specific antibodies against different subtypes of influenza virus. Also, it is affordable for seroepidemiology in poor and limited resource settings. Co-infection of HIV with influenza has been identified as one of the causes of severe respiratory diseases in high HIV endemic settings including sub-Saharan Africa. In South Africa, 44 % of influenza positive patients having acute respiratory infections are HIV infected (10).

This hospital population-based study revealed an overall 39.7 % (69/174) serological prevalence of influenza among HIV positive patients on cART. The prevalence rate is comparable with the 31 % serological response recorded in HIV positive individuals in Miami, USA (18). A significantly lower seroprevalence rate of 14.7 % was recorded in Taiwan (19). A higher incidence rate of 71.6 % in HIV patients was recorded in Rome, Italy by Agrati, Gioia (20). In Africa, studies done by Cohen, Walaza (21) reported seroprevalence of 29.8 % in South Africa. Ho, Aston (22) recorded a seroprevalence of 25.4 % in Malawi, similar to that of Ope, Katz (23) that reported an incidence rate of 24.5 % in Kenya. This could be attributed to differences in a lot of factors including sample size and assay type used.

Our investigation further revealed that 84.1 % (58/69) were IgM seropositive for influenza A while 15.9 % (11/69) were positive for influenza B virus, signifying epidemiological dominance of influenza A over influenza B virus in the third quarter of 2018 in Lagos, Nigeria. Ho, Aston (22) reported that conversely in low resource settings with high HIV prevalence, there is high incidence of influenza illness and greater risk of hospitalization among HIV infected person which explains the high prevalence rate of influenza A detected in this study. A study in Germany recorded higher seroprevalences of 96.3 % for influenza A and 98 % for influenza B using ELISA method, although the study measured IgG antibody in non-HIV patients (17). The higher incidence could be attributed to the fact that IgG antibody indicates past infection which may sometimes include previous vaccination, hence the finding is usually significantly higher than a measure of IgM antibody. Nonetheless, the current study population had never been previously vaccinated due to unavailability of influenza vaccine in Nigeria.

Interestingly, 13.4 % (9/69) of the HIV patients had seropositivity to both influenza A and B viruses. Coinfection of influenza A and B viruses in HIV patients may exacerbate the immune status by impairing T-cell response with resultant fatal consequences in such individuals. It is evident in previous reports that influenzaA(H1N1)pdm09 (now seasonal influenza virus) caused severe pneumonia leading to acute respiratory distress syndrome (ARDS) and multiple organ dysfunction associated with death rate of 17‒ 54 % in HIV patients (24, 25). Hence, this calls for serious public health concern and immediate action at ameliorating the menace of influenza co-infection in people living with HIV (PLWHIV).

Our study demonstrated the highest 39.1 % seroprevalence of influenza among HIV patients in the age group 41‒ 50 years. On the contrary, Tempia, Walaza (26) recorded the highest seroprevalence among age group 25‒ 44 years. Furthermore, our finding disagrees with the CDC report that persons above 65 years are more likely to be at risk for influenza virus (27). This is however supported by Lambert, Ovsyannikova (28) that on the account of hospitalization and mortality rates, older adults do not have the highest rate of infection and does not represent main contributors to local outbreaks. This may also be due to rapid waning of immunity among the elders (17).

Based on gender, a higher 65.2 % seroprevalence was detected among females compared to the males with 24.6 %, a finding that is consistent with that of Ho, Mallewa (29) that reported higher seroprevalence among the females. On the contrary, Patel, Bush (9) recorded a higher seroprevalence in males than the females. A suggestive reason for the higher seroprevalence among females could be the overwhelming burden of HIV that affects more women in sub-Saharan Africa making them to be more susceptible to influenza infection.

Out of the 19/69 (27.5 %) immunocompromised patients, 4/19 (21.1 %) were severely immunosuppressed (CD4 < 200 cells/mm^3^). This is in line with the study of Tempia, Walaza (26) that reported a seroprevalence of 22.6 % in immunocompromised patients. However, the role of CD4 and their impact on HIV positive patients is less well understood partly due to their heterogeneity and lack of epitope specific systems (30). Association between the incidence of influenza infection and CD4 count showed that HIV patients with CD4 < 400 had significantly higher prevalence. Contrary to the study of Cohen, Moyes (11) and Tempia, Walaza (26), the prevalence of influenza was higher among individuals with CD4 count > 400 compared to those with lower CD4 count. Similar to this study, Patel, Bush (9) recorded higher seroprevalence in patients with CD4 count > 200 cells/mm^3^ than those < 200 cells/mm^3^. A good reason for these observations can be explained based on the traditionally accepted role of influenza specific CD4 T cells in providing help to B cells for the production of high-quality antibodies. Hence, depletion of CD4 T cells prior to influenza challenge results in dramatic drop of antibodies’ titers.

People with viral load < 100 copies/mL (73.9 %) had the highest seroprevalence of influenza virus. This agrees with the work of Patel, Bush (9) that recorded seroprevalence of 85 % for patients with viral load < 400 copies/mL. This also indicates that reduced viraemia of HIV doesn’t reduce the susceptibility of HIV patients to influenza virus.

Interestingly, one of the patients seropositive for influenza B was positive for HBsAg and had a CD4 count of 162 while another patient positive for HBsAg and influenza A had a CD4 count of 570 which doesn’t really explain the particular CD4 range where influenza might become a problem, but showed comorbidity that can lead to fatal consequence in such patients.

Among the 165 HIV patients on fixed dose combination (FDC) of some regularly prescribed cART including Tenofovir, Emtricitabine, Azidothymidine, Atazanavir, Efavirenz, Nevirapine, Lopinavir/ Ritonavir, 61 were seropositive for influenza A and B viruses. The direct relationship of cART on patients and their seropositivity to influenza is not well understood. Nonetheless, our study demonstrated that HIV patients on cART were susceptible to recent infection of influenza viruses. In support of this finding, an ecological study done by Cohen, McMorrow (2) estimated an increased risk of influenza associated mortality among HIV positive individuals even after the widespread introduction of cART. Also, a study by Parmigiani, Alcaide (18) concluded that regardless of vaccination and control of viral load with cART, HIV infected patients are at an elevated risk of acquiring seasonal influenza infection. On the contrary, Jambo, Sepako (31), opined that in advanced countries, highly active antiretroviral therapy (HAART) introduction has led to the reduction of influenza related complications among HIV infected patients.

Part of the limitations of this study is that the study failed to consider other risk factors for influenza and HIV co-infection such as smoking, and intravenous drug use. Also, serological evidences from different HIV treatment centres are necessary for representative data in Nigeria.

## Conclusions

This study demonstrated that HIV patients are susceptible to influenza both Influenza A and B viruses. It however showed low seroprevalence of coinfection of influenza A and B and that co-infection may further impair the immunocompromised system of HIV patients. It has also provided information on the impact of CD4 cell count and cART on influenza incidence in highly suppressed HIV RNA patients. Nonetheless, co-infection with HbsAg and an immunosuppressed state may further predispose them to serious influenza life threatening complications. Strong advocacy on the need to reduce risk of exposure to influenza, and provision of influenza vaccine in Nigeria are recommended to prevent such complications.

## Acknowledgements

We sincerely acknowledge all the patients that volunteered to be part of this study. We acknowledge the LASUTH Directorate of Clinical services and Training / Hospital Research Ethics Committee for the approval of the study research proposal. We are grateful to the Management and Staff of APIN- LUTH for the approval for sample collection. Special thanks to all the Laboratory Technologists that supported the student’s research work. Many thanks to all the senior Faculty members that criticized and reviewed the initial manuscript. The abstract for this study was accepted for presentation at the 52^nd^ Miami Winter Symposium 2019: Evolving Concepts in HIV & Emerging Viral Infections Organized by Elsevier, University of Miami Miller School of Medicine, International Union of Biochemistry & Molecular Biology in Florida, USA (January 27-30^th^, 2019).

